# Microbial associations and spatial proximity predict North American moose (*Alces alces*) gastrointestinal community composition

**DOI:** 10.1101/514604

**Authors:** Nicholas M. Fountain-Jones, Nicholas J. Clark, Amy C. Kinsley, Michelle Carstensen, James Forester, Timothy J. Johnson, Elizabeth Miller, Seth Moore, Tiffany M. Wolf, Meggan E. Craft

**Author notes:** Department of Veterinary Population Medicine, University of Minnesota, 1365 Gortner Avenue, St Paul, Minnesota 55108 (Nicholas M. Fountain-Jones).

## Abstract

1. Microbial communities are increasingly recognised as crucial for animal health. However, our understanding of how microbial communities are structured across wildlife populations is poor. Mechanisms such as interspecific associations are important in structuring free-living communities, but we still lack an understanding of how important interspecific associations are in structuring gut microbial communities in comparison to other factors such as host characteristics or spatial proximity of hosts.
2. Here we ask how gut microbial communities are structured in a population of North American moose (*Alces alces*). We identify key microbial interspecific associations within the moose gut and quantify how important they are relative to key host characteristics, such as body condition, for predicting microbial community composition.
3. We sampled gut microbial communities from 55 moose in a population experiencing decline due to a myriad of factors, including pathogens and malnutrition. We examined microbial community dynamics in this population utilizing novel graphical network models that can explicitly incorporate spatial information.
4. We found that interspecific associations were the most important mechanism structuring gut microbial communities in moose and detected both positive and negative associations. Models only accounting for associations between microbes had higher predictive value compared to models including moose sex, evidence of previous pathogen exposure, or body condition. Adding spatial information on moose location further strengthened our model and allowed us to predict microbe occurrences with ∼90% accuracy.
5. Collectively, our results suggest that microbial interspecific associations coupled with host spatial proximity are vital in shaping infra communities in a large herbivore. In this case, previous pathogen exposure and moose body condition were not as important in predicting gut microbial community composition. The approach applied here can be used to quantify interspecific associations and gain a more nuanced understanding of the spatial and host factors shaping microbial communities in non-model hosts.

## Introduction

The relative roles of interspecific associations versus the environment in shaping communities are intensely debated (e.g., Chase & Myers, 2011; Vellend et al., 2014). In particular, detecting negative (e.g., potential competition) or positive (e.g., potential facilitation) associations between organisms remains a fundamental challenge of community ecology (e.g., Barner, Coblentz, Hacker, & Menge, 2018; Dormann et al., 2018; Harris, 2016; Ovaskainen et al., 2017). Despite the challenges of detecting associations between co-occurring species, associations between species are thought to be essential for structuring free-living communities (e.g., de Araújo, Marcondes-Machado, & Costa, 2014). For example, associations between forest animal communities were more important in structuring communities than habitat attributes such as vegetation (Le Borgne et al., 2018). In contrast, the roles that interspecific associations play in governing the assembly of microbial systems (*microbial interspecific associations*) are less understood (Ganz et al., 2017; Herren & McMahon, 2018; Zelezniak et al., 2015), particularly for within-host microbial communities. Even when microbial interspecific associations are quantified in infra-communities, the relative importance of microbe dispersal (Evans, Martiny, & Allison, 2017; Zhou & Ning, 2017) and host characteristics in shaping microbial infra-communities is rarely assessed (Clark, Wells, & Lindberg, 2018b). The quantification of interspecific associations of gut microbes is rare in wildlife populations but, given the significant ecological insights derived from studies of human gut microbial communities (e.g., Faust et al., 2012), is an important knowledge gap to fill.

While our understanding of how microbial communities are structured is poor, it remains a key goal in microbial ecology, particularly given microbial community relevance to animal health. Host characteristics such as sex, body condition, and the presence or absence of pathogens are recognised as important for shaping microbial infra-communities (Britton & Young, 2014; Cheng et al., 2015; Ganz et al., 2017; Hooper, Littman, & Macpherson, 2012; Jani & Briggs, 2014, 2018; McKenney et al., 2015; Mshelia et al., 2018; Näpflin & Schmid-Hempel, 2018). For example, horses in poor body condition have greater microbial diversity and a different suite of microbial species present compared to horses in good body condition (Mshelia et al., 2018). High loads of the pathogen *Batrachochytrium dendrobatidis* can not only increase amphibian skin microbial diversity but also alter microbial composition (Jani & Briggs, 2014, 2018). However, the relative importance of host characteristics compared to other ecological processes, such as interspecific associations, in structuring microbial infra-communities is poorly understood.

Understanding how microbial infra-communities are shaped, however, is technically challenging, in part due to the complexity of untangling complex ecological processes in often species-rich but poorly characterized communities (Zhou & Ning, 2017). Graphical network models such as conditional random fields (CRF) offer exciting opportunities to address this challenge by estimating associations between organisms in a community (Clark, Wells, & Lindberg, 2018b). In effect, these models can untangle relative influences of microbial interspecific associations and host characteristics in predicting community compositions. Crucially, inferences gleaned from CRFs can be integrated with phylogenetic or functional data to provide new insights into mechanisms underlying microbial community assembly. For example, associations between microbes may be non-random in that phylogenetically or functionally similar species may co-occur more frequently (e.g., Bauer & Thiele, 2018). However, non-random associations detected in the human gut have been found to be often negative and between phylogenetically or functionally distinct bacterial species, indicating that competition is likely an important driver of structure in this community (Faust et al., 2012). Overlaying information such as this on graphical models has the potential to highlight the broader mechanisms shaping microbial infra-communities.

Here we use a CRF approach to investigate microbial community composition in a wild moose (*Alces alces*) population in Minnesota. Over the last two decades, moose in Minnesota, which exist on the southern edge of their species range, have experienced significant population decreases (Delgiudice, 2018; Lenarz, 2009). The most dramatic decline has been reported for moose in northwest Minnesota, where the population declined from 4,000 animals in the 1980s to less than 100 by the mid-2000’s due to a combination of pathogens and malnutrition (Murray et al., 2006). Recently, moose in northeast Minnesota have experienced a 55% population decline, driven largely by parasitic infections in adults and wolf predation in calves (Carstensen et al., 2018; Severud, 2017; Wünschmann et al., 2015). Moose gut microbial communities are known to vary with sex and age (Ishaq, Sundset, Crouse, & Wright, 2015; Ishaq & Wright, 2012), but to what extent these host characteristics, as well as pathogens and malnutrition, shape moose gut microbial communities is unknown. Other ecological processes such interspecific assocations, even though rarely quantified in wild populations, are also likely to play a role in shaping the moose gut microbial community as they do for other ruminants (Henderson et al., 2015). Furthermore, animals within close proximity to each other may have similar gut microbial communities due to similarities in diet and/or increased possibilities for microbial dispersal between individuals; both factors known to be important in gut microbial community assembly in many host species (Henderson et al., 2015; Moeller et al., 2017).

Here we apply a novel CRF approach that can quantify the relative importance of host characteristics (including moose sex, body condition, pathogen exposure, and pregnancy status) and microbial interspecific associations in shaping moose gut communities whilst accounting for underlying spatial autocorrelation in microbial occurrences. Accounting for spatial autocorrelation not only reduces false detection of interspecific associations that may arise due to dispersal limitation or diet but quantifies how important these processes could be in shaping microbial distributions. Specifically, we address the following questions:

1. Do CRF model combinations including host characteristics and spatial proximity outperform models reflecting just associations between microbes in explaining microbial communities?
2. After controlling for host characteristics and spatial patterns are there any remaining non-random negative or positive associations between microbes?
3. If there are non-random associations, are they between microbes that are functionally or phylogenetically similar?

We hypothesized that host characteristics would be the dominant processes shaping the moose gut microbial community, and that spatial proximity between hosts and interspecific associations would also partially explain gut microbial co-occurrence.

## Materials and Methods

### Sample collection and sequencing

Faecal samples were collected from 55 wild moose that were part of companion studies of survival and cause-specific mortality led by the Minnesota Department of Natural Resources, Grand Portage Band of Lake Superior Chippewa, and Voyageur’s National Park across north-eastern Minnesota (Fig. 1), placed on ice until transportation to the laboratory, and stored at - 20°C prior to processing. This included moose that were live-captured (n = 52, 2011-2015) and sick (n = 3, 2009-2010). Details of the capture and handling of these moose can be found in Butler et al. (2012) and Carstensen et al. (2018). The metadata provided from these moose included date and location of capture, pregnancy status, sex, age, body condition, and serological exposure to *Borrelia burgdorferi*, West Nile Virus, and leptospira (6 serovars including *L. bratislava, L. canicola, L. grippotyphosa, L. hardjo, L. icterohemorrhagica* and *L. pomona*). In total, we had serological data for 49 moose (33 females and 16 males) with the remaining 6 individuals removed from the CRF analysis (but retained for the descriptive analysis). See Butler et al. (2012) for serological test details. Of the live-captured moose, 21 were observed to be underweight (thin/very thin), and 31 were considered of normal body condition at winter capture (Jan-Feb). All samples apart from one were collected in winter between 2011-2015; the other sample was collected in 2003. Microbial DNA was extracted from the faecal samples using the PowerSoil DNA Isolation kit (Qiagen), in accordance with the manufacturer’s protocol. The V4 hypervariable region of the bacterial 16S rRNA gene was amplified using the barcoding primers 515F and 806R. Amplicons were sequenced on the Illumina MiSeq platform following the method outlined by Gohl et al., (2016).

**Figure 1:**
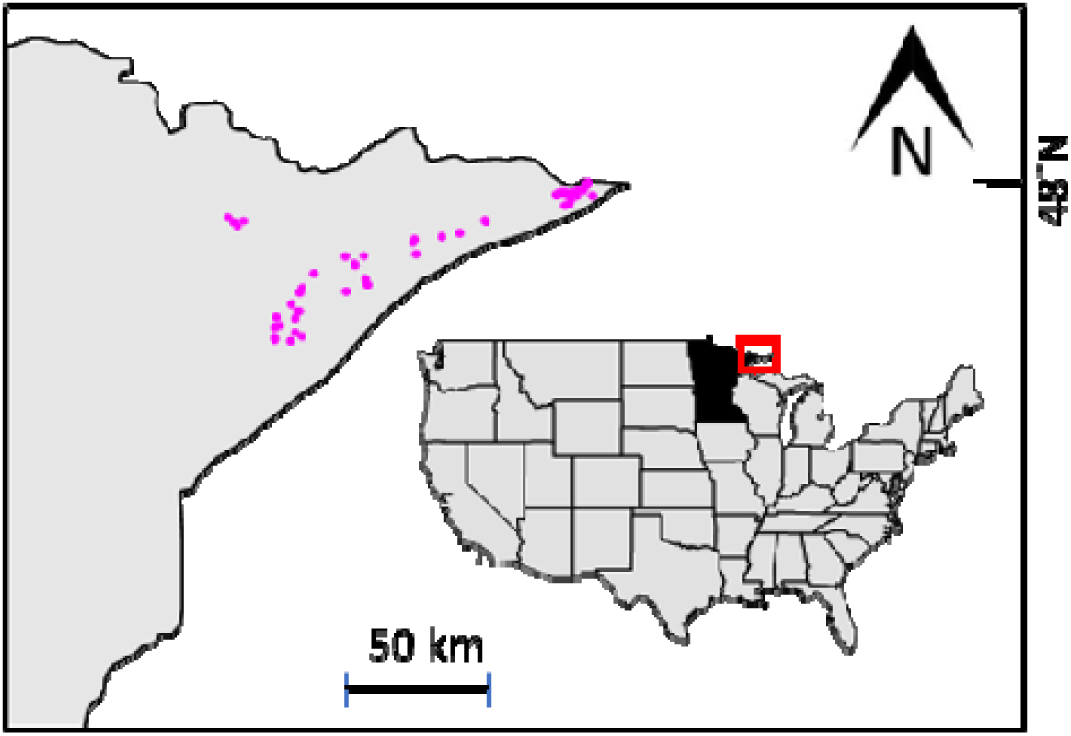
Locations (pink dots) of moose faecal samples in northeastern Minnesota, USA. The red box shows the location of the study in Minnesota.

### Bioinformatics pipeline

Raw sequencing reads were processed using the University of Minnesota’s metagenomics-pipeline (complete description of the pipeline can be found at https://bitbucket.org/jgarbe/gopher-pipelines/overview), which implements the QIIME version 1.9.1 analysis software (Caporaso et al., 2010). Briefly, sequencing adaptors were trimmed using Trimmomatic (Bolger, Lohse, & Usadel, 2014) and read pairs were assembled and primer sequences removed using PANDAseq (Masella, Bartram, Truszkowski, Brown, & Neufeld, 2012). Reads without primers, unpaired reads, and assembled reads that were outside the expected rRNA gene V4 region length were discarded. Chimeras were detected and removed with QIIME’s identify_chimeric_seqs.py function with the usearch61 algorithm (Edgar, 2010). Open-reference OTU (operational taxonomic unit) picking was conducted using QIIME’s pick_open_reference_otus.py with a minimum sequence identity threshold of 97%. Representative OTU sequences were aligned against the Greengenes 18.8 core set (DeSantis et al., 2006b) using UCLUST (Edgar, 2010) with QIIME default parameters. Singleton OTUs and those that did not align with PyNAST (Caporaso et al., 2010) were removed from the analysis. To control for differences in sequencing depth between samples, read counts were rarefied to the lowest number of reads (101,131) per sample.

To test for co-occurrence patterns between gut microbes, we filtered out OTUs with ≤ 10% abundance (i.e., only keeping OTUs that occurred in at least 10% of samples) and converted the OTU table into a presence-absence matrix. OTUs that were found ≥ 75% of samples were also filtered from the analysis. Our purpose for removing rare and common OTUs was to ensure that adequate inferences could be made about the occurrence probability of each OTU (Ovaskainen, Abrego, Halme, & Dunson, 2016). This would be difficult / impossible to do if the target OTU shows little variability across sampled environmental gradients by being either too rare or too common. We transformed the presence-absence matrix into a Jaccard similarity matrix and used non-metric multidimensional scaling (NMDS) to select OTUs that were driving compositional change across the moose samples and therefore likely to differ with moose characteristics such as body condition (Pearson correlation coefficient > 0.45). In total, 42 OTUs remained after this filtering process. NMDS was performed using functions in the R package *vegan* (Oksanen et al., 2013).

To reduce the dimensions of the serological exposure data for subsequent analysis, we converted test results into a binary matrix (0 if the individual tested negative and 1 if positive), transformed the binary matrix into a Jaccard similarity matrix, and then performed principal coordinate analysis (PCoA) using the R package *vegan* (Oksanen et al., 2013). The first three PCoA axes represented 85% of the variation in exposure across individuals and eigenvalues from these axes were used as covariates in the CRF analysis below (now called *pathogen exposure PCoA axes*).

### Conditional random fields

#### Identifying OTU co-occurrence patterns using conditional random fields

The framework we used to investigate OTU co-occurrence probabilities while accounting for potential influences of covariates is described in detail by Clark et al. (2018a) and references therein. Briefly, the log-odds of observing OTU *j* given covariate *x* and the presence-absence of OTU *k* is modelled using:

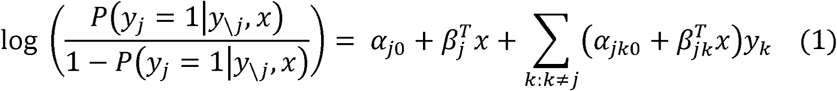

where *y*_*j*_ is a vector of binary observations for OTU *j* (1 if the OTU was present, 0 if absent), *y*_\*j*_ represents vectors of binary observations for all other OTUs apart from *j, α*_*j0*_ is the OTU-level intercept, and *β*^*T*^_*j*_ is the coefficient for the effect of covariate *x* on OTU *j*’s occurrence probability. Interaction parameters are represented by *α*_*jk*0_ and 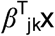 (defined below). Parameterization of the likelihood is estimated using logistic regression, where regression coefficients represent the effects of predictors on the OTU’s conditional log-odds. Cross-multiplying all combinations of co-occurring OTUs and external covariates allows direct comparison of the relative influences interspecific associations) and host effects on an OTU’s occurrence probability. For each OTU-specific regression, sparsity is added to the model using L1 (e.g., least absolute shrinkage and selection operator [LASSO]) regularisation to force regression coefficients toward zero if they have minimal effects. Ten-fold cross-validation was implemented to choose the penalty that minimised cross-validated error, as this is considered an appropriate loss function in binomial classification studies. Following LASSO variable selection, coefficients representing conditional dependence of two OTUs and coefficients representing effects of covariates on this dependence were symmetrized by taking the mean of the corresponding estimates so that *α*_*jk0*_ = *α*_*kj0*_ and 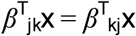. This means we can approximate parameters from a unified graphical network, after maximizing the conditional log-likelihood for each OTU (Lee & Hastie, 2015). Following unification, inference is straightforward. If *α*_*jk0*_ = 0, we can infer that the occurrence probabilities of OTUs *j* and *k* are conditionally independent, after accounting for covariates and other OTUs. If *α*_*jk0*_ ≠ 0 but 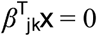, the occurrence probabilities are still considered conditionally dependent, but the strength of this dependence is not expected to co-vary with covariate *x*.

#### MRF and CRF model selection

We estimated four graphical model formulations of increasing complexity to identify a best-fitting model for our OTU presence-absence dataset. In the first, we built a Markov random fields (MRF) graph that did not include any spatial data or host characteristics and used only the binary occurrences of the 42 OTUs as predictors (hereafter the *MRF model*). In the second model, the GPS coordinates for each observation (latitude and longitude, in decimal degrees) were used to construct penalised Gaussian process regression splines with 100 degrees of freedom (Kammann & Wand, 2003; Wood, 2003) (hereafter the *spatial MRF model*). Including the spatial splines in each OTU’s linear predictor ensured that interspecific associations on occurrence probabilities were estimated only after accounting for possible spatial autocorrelation. For the third model, we built a CRF including moose characteristics covariates (moose sex, body condition, pregnancy status and the pathogen exposure PCoA axes) as well as longitude and latitude (hereafter the *non-spatial CRF model*). All covariates were included as scaled continuous variables with the exceptions of sex and pregnancy status, which were both included as categorical variables. For the final model, we built a CRF as above, but we replaced the latitude and longitude variables with spatial splines (hereafter the *spatial CRF model*).

We assessed the fit of each of the above candidate models to the observed data by calculating the proportion of observations that each model successfully classified. This was done using ten-fold cross-validation. The best-fitting model was then fit to 100 bootstrapped versions of the observed data (randomly shuffling observations in each bootstrap iteration) to capture uncertainty in model parameters. All CRF model fitting was performed using functions in the *MRFcov* R package (Clark, Wells, & Lindberg, 2018a). From the OTU co-occurrence data, we constructed an adjacency matrix and plotted association networks using *iGraph* R package (Csárdi & Nepusz, 2006). See Appendix S1 for R code detailing data preparation, analytical routine and model specification.

## OTU functional predictions

We predicted the molecular functions of OTUs using PICRUSt’s precalculated functional prediction table, where the rows were KEGG orthologs (KOs) and the columns were OTUs based on Greengeens identification numbers (DeSantis et al., 2006a; Kanehisa, Sato, Kawashima, Furumichi, & Tanabe, 2016; Langille et al., 2013). This KO table was converted into a presence-absence matrix (i.e., whether a particular functional ortholog is associated with that OTU) and calculated the pairwise functional similarity using the Jaccard index. We applied NMDS to view the broad functional relationships between OTUs and define functional groups (FGs).

## Results

We found that Firmicutes, followed by Bacteroidetes, were the dominant phyla in all our moose gut communities making up over 75% of the reads detected (Fig. S1). The ratio of these groups across samples was relatively consistent with only one individual having a proportion < 50% of Firmicutes present (Fig. S2). Functionally the OTUs could be grouped into two FGs (Fig. S3).

### Model fit

The MRF model, which only quantified associations between microbes, could more accurately predict OTU occurrence compared to either the spatial CRF or non-spatial CRF models (both of which included host characteristics; Fig. 2a). Including spatial regression splines did improve the fit of the CRF model, with this model accurately predicting 78% of observations compared to 74% of for the non-spatial CRF (Fig. 2a). In contrast, the MRF could accurately predict 83% of presence-absence observations on average (Fig. 2a). This result was not due to one model being better able to predict occurrence over absence, as the MRF had much higher specificity and sensitivity than either of the CRF models (Fig. 2b/c). However, we still found evidence of spatial clustering in microbial occurrence probabilities, as MRF performance improved further when we included spatial splines in the model (Fig. 2a). Adding spatial data to this model enabled us to correctly predict ∼90% of OTU occurrences in the moose gut microbial community. This not only suggests that microbes in our dataset were more likely to show similar occurrences in moose that were sampled nearby to one another, but that false associations between microbes could be inferred by models that do not account for spatial proximity.

**Figure 2:**
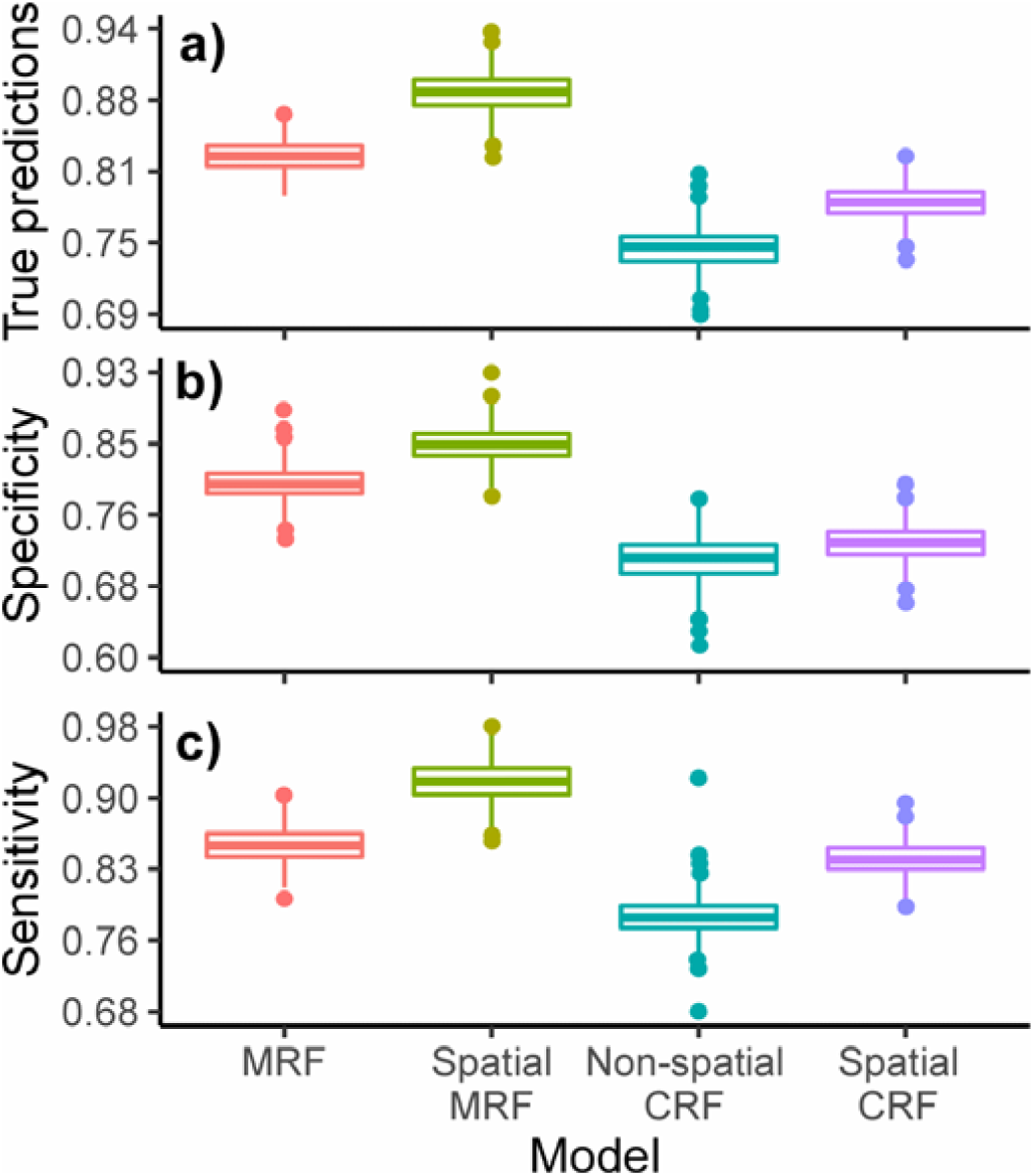
Box and whisker plots showing the predictive performance of the Markov random field model (MRF – without model covariates), spatial MRF model, non-spatial conditional random field model (CRF - with model covariates) and spatial CRF as defined by (a) true prediction performance, b) specificity, and c) sensitivity. Predictions were assessed by fitting models to a random fold containing 90% of the data (training data) and predicting observations for the remaining 10% (test data). This process was repeated 100 times to capture uncertainty in performance. Specificity is the ability of the model to correctly identify individual moose without the specified OTU (proportion of observed negatives that were predicted to be negative), while sensitivity is the ability to correctly identify individuals with the specified OTU (proportion of observed positives that were predicted to be positive). Box hinges show the interquartile range (25% and 75% quantiles), lines within boxes indicate the median (50% quantile), whiskers show 10% and 90% quantiles and dots show values outside these quantiles.

### Microbial associations

The co-occurrence network revealed that both positive and negative associations occurred across taxonomic and functional groups with no clear phylogenetic pattern (i.e., OTUs from the same phyla/class or function were not preferentially negatively or positively associated with each other, Fig. 3). On average, we found that positive associations between OTUs were more common than negative associations (4 vs 2, Table 1). Firmicutes was the best-represented phyla in our moose faecal samples (Fig. S1), and this phylum also had the highest number of OTU associations. Strikingly, even though Bacteroidetes was the second most dominant phyla, we did not detect any associations involving OTUs from this taxa. In contrast, OTUs from Tenericutes were rare (3% of sequences, Fig. S1), but overall, they had on average 1 more positive association partner than other phyla (4 vs 3), whereas Cyanobacteria OTUs had more negative association partners than other phyla (3 vs 2, Table 1). FGs showed similar differences, with FG1 (which was dominated by the Tenericutes; Fig. S3) having more positive associations compared to FG2 (4 vs 3 on average). The opposite was true for negative associations, with FG2 having more associations on average than FG1 (1 vs 2). Overall, two previously uncharacterized OTUs (identified by the ‘NewReference’ label) from class Mollicutes (Tenericutes) and class Clostridia (Firmicutes) had the highest number of associations overall (8, Table 1). Two OTUs from class Clostridia had the highest numbers of negative associations (5 & 6 respectively, Table 1).

**Table 1:** Summary of associations detected in the MRF analysis.

| OTU identity | Avg.oc | Pos | Neg | Phylum\|Class | FG |
| --- | --- | --- | --- | --- | --- |
| 338145 | 0.2043 | 6 | 4 | Actinobacteria\|Coriobacteriia | 2 |
| NewReferenceOTU314 | 0.6637 | 2 | 1 | Actinobacteria\|Coriobacteriia | N |
| NewReferenceOTU69 | 0.6832 | 2 | 1 | Actinobacteria\|Coriobacteriia | N |
| NewReferenceOTU74 | 0.7562 | 3 | 2 | Actinobacteria\|Coriobacteriia | N |
| 206930 | 0.4799 | 4 | 4 | Cyanobacteria\|4C0d-2 | 2 |
| NewReferenceOTU123 | 0.5473 | 6 | 3 | Cyanobacteria\|4C0d-2 | N |
| NewReferenceOTU73 | 0.5461 | 0 | 2 | Cyanobacteria\|Chloroplast | N |
| 322906 | 0.9437 | 3 | 4 | Firmicutes\|Clostridia | 2 |
| 325706 | 0.6148 | 0 | 1 | Firmicutes\|Clostridia | 2 |
| 4317006 | 0.0372 | 5 | 4 | Firmicutes\|Clostridia | 2 |
| 4480841 | 0.5291 | 2 | 1 | Firmicutes\|Clostridia | 2 |
| 4314603 | 0.4536 | 2 | 1 | Firmicutes\|Clostridia | 2 |
| 4417708 | 0.0843 | 7 | 4 | Firmicutes\|Clostridia | 2 |
| 296918 | 0.7692 | 1 | 4 | Firmicutes\|Clostridia | 2 |
| 266952 | 0.6309 | 3 | 3 | Firmicutes\|Clostridia | 2 |
| 294064 | 0.5084 | 5 | 3 | Firmicutes\|Clostridia | 2 |
| 294262 | 0.2403 | 6 | 1 | Firmicutes\|Clostridia | 2 |
| 579159 | 0.2068 | 3 | 2 | Firmicutes\|Clostridia | 2 |
| 333577 | 0.3276 | 2 | 1 | Firmicutes\|Clostridia | 2 |
| 1038874 | 0.3899 | 2 | 0 | Firmicutes\|Clostridia | 2 |
| 129755 | 0.3879 | 3 | 0 | Firmicutes\|Clostridia | 2 |
| 738351 | 0.6364 | 1 | 0 | Firmicutes\|Clostridia | 2 |
| 4295783 | 0.3693 | 1 | 0 | Firmicutes\|Clostridia | 2 |
| 574585 | 0.6326 | 1 | 5 | Firmicutes\|Clostridia | 2 |
| NewReferenceOTU108 | 0.0809 | 8 | 3 | Firmicutes\|Clostridia | N |
| NewReferenceOTU74 | 0.3172 | 1 | 0 | Firmicutes\|Clostridia | N |
| NewReferenceOTU76 | 0.2891 | 5 | 6 | Firmicutes\|Clostridia | N |
| NewReferenceOTU76 | 0.0827 | 6 | 4 | Firmicutes\|Clostridia | N |
| NewReferenceOTU157 | 0.0899 | 3 | 0 | Firmicutes\|Clostridia | N |
| NewReferenceOTU289 | 0.1515 | 5 | 4 | Firmicutes\|Clostridia | N |
| 4396877 | 0.0928 | 5 | 1 | Firmicutes\|Erysipelotrichi | 2 |
| NewReferenceOTU42 | 0.5286 | 3 | 4 | Firmicutes\|Erysipelotrichi | N |
| 339838 | 0.063 | 1 | 1 | Tenericutes\|Mollicutes | 1 |
| 513605 | 0.0299 | 7 | 1 | Tenericutes\|Mollicutes | 1 |
| 4446732 | 0.6627 | 4 | 2 | Tenericutes\|Mollicutes | 1 |
| 1108356 | 0.5123 | 4 | 2 | Tenericutes\|Mollicutes | 1 |
| 921020 | 0.1596 | 5 | 1 | Tenericutes\|Mollicutes | 1 |
| NewReferenceOTU413 | 0.2271 | 5 | 2 | Tenericutes\|Mollicutes | N |
| NewReferenceOTU493 | 0.2445 | 4 | 1 | Tenericutes\|Mollicutes | N |
| NewReferenceOTU390 | 0.6923 | 0 | 0 | Tenericutes\|Mollicutes | N |
| NewReferenceOTU404 | 0.1345 | 8 | 2 | Tenericutes\|Mollicutes | N |
| NewReferenceOTU528 | 0.4842 | 4 | 4 | Tenericutes\|RF3 | N |
| Average |  | 4 | 2 |  |  |

Avg.oc: average occurrence in our moose samples. Pos: number of positive associations.
Neg: number of negative associations. 'NewReference' indicates that the OTU has not been
previously characterised by the Greengenes database. FG: Functional group (see Figure S3).
N: No functional group could be determined (not previously characterized).

**Figure 3:**
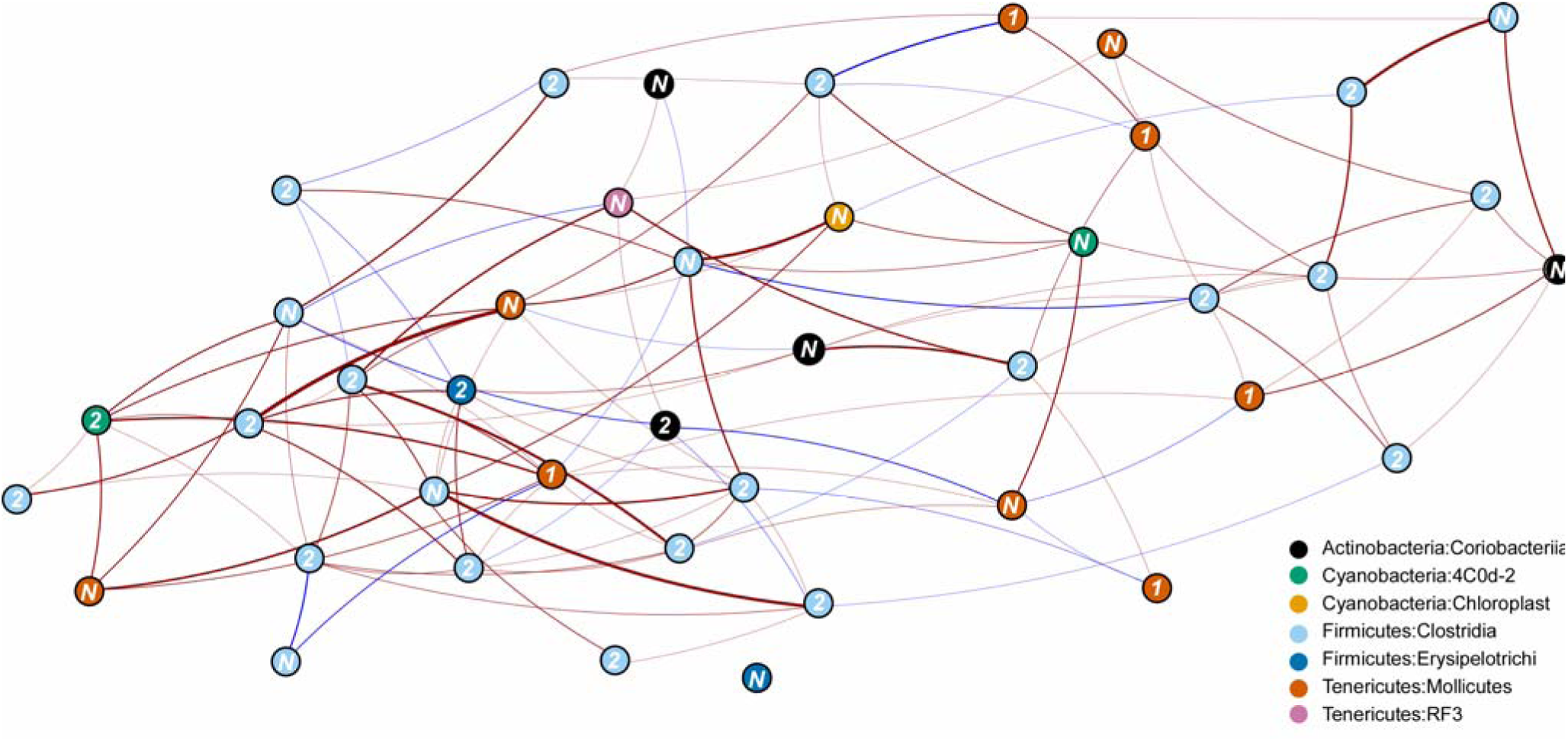
Moose gut microbial MRF co-occurrence network. Blue edges indicate negative associations and red edges represent positive associations. Edge thickness is scaled by the strength of association. Node colour indicates taxonomic group of each microbe. Numbers on the node represent which broad functional group (FG) each OTU belonged to (see Fig. S3). ‘N’ indicates that there was no functional data for this microbe. See Fig. S4 for the OTU correlation matrix with names included.

## Discussion

Utilizing graphical network models, we show the importance of microbial interspecific associations over host characteristics in shaping moose gut microbial communities at a population scale. In this case, both interspecific associations and spatial proximity were important for shaping microbial communities in this declining moose population. Host characteristics were relatively less important in predicting the distribution of microbes. Even though we did not have host genetic or specific diet data from individuals (i.e. stable isotope data), we could predict microbial distributions using just interspecific associations with remarkably high accuracy. Across this moose population, we detected non-random negative and positive microbial associations with no clear functional or phylogenetic pattern. Our study not only highlights the importance of accounting for interspecific associations when trying to quantify how host characteristics shapes host infra-communities but also shows the value of graphical network models in untangling community dynamics more broadly.

We found that evidence of pathogen exposure was not particularly important in predicting moose gut microbial community dynamics. As the serological evidence we used in this study only infers past infection by a particular pathogen rather than current infection status, perhaps this is not surprising. Previous studies testing the role of pathogens shaping microbial communities have used evidence of current infection (e.g., qPCR; Ganz et al., 2017; McKenney et al., 2015) rather than previous exposure based on serological evidence. When we sampled the moose gut microbial communities, the animal may not be experiencing the infection, and this may be the reason we did not detect an effect. It is also possible that herbivore gut microbial communities are more resilient to pathogen infection than other trophic groups such as carnivores. Ruminant microbial communities are more dominated by environmental bacteria cultivated by the host to aid digestion (Muegge et al., 2011) and may be less regulated by the immune system compared to the gut microbial communities in other species (Ley et al., 2008). Lower immune regulation, for example, could explain why the herbivores such as the gorilla (*Gorilla gorilla*) have microbiomes remarkably resilient to Simian immunodeficiency virus infection compared to omnivorous chimpanzees (*Pan troglodytes*) or for HIV infection in humans (Moeller et al., 2015). In support of this, we find that across individual moose in our study, the relative proportions of gut microbial taxa were relatively consistent, despite large differences in body condition and exposure status (Fig. S2). Similar ratios of the different microbial taxa in moose have also been reported in moose from the east coast of North America (Ishaq & Wright, 2012) even though samples were collected from moose in a different season. Further studies where current pathogen infection status is known in moose and other herbivores are needed to resolve this question.

Interspecific associations, in contrast, were much better predictors of moose gut microbial communities. In free-living communities, the role of interspecific associations in shaping the distributions of species is increasingly recognised (e.g., Aragón, Carrascal, & Palomino, 2018; C. B. de Araújo et al., 2014). However, the importance of interspecific associations is likely to decrease with geographic scale (Araújo & Rozenfeld, 2014). Whether this applies to gut microbial communities, in contrast, is poorly studied. In our case, host effects may have been detectable if we sampled a greater number of individuals across a larger portion of moose home range in central North America. Nonetheless, the scale at which wildlife surveillance is performed (and thus faecal samples collected for gut microbiome analysis) is often similar across studies (e.g., Cheng et al., 2015). In general, understanding the role that scale plays in structuring gut microbial communities, beyond the population scale, is a major challenge for microbial ecology (Antwis et al., 2017; Camp, Kanther, Semova, & Rawls, 2009). Based on our findings, we suggest that larger geographical and/or temporal scales are needed to detect possible impacts of host characteristics on the structure of gut microbial communities.

Even without accounting for host traits such as host genetics that are thought to be important in shaping the gut microbial communities in other herbivores (Kohl, Varner, Wilkening, & Dearing, 2017), our models had high predictive performance. Our results may be due to samples coming from a relatively well-mixed population, even though our moose samples were taken over a large area spanning a maximum of ∼150 km. Central North America (including Minnesota) has the highest genetic diversity of all North American moose populations (Hundertmark, Bowyer, Shields, & Schwartz, 2003), but to what extent our samples come from genetically distinct populations is unknown. Future work incorporating host genetics in moose may help better understand the role of host characteristics, compared to microbial interspecific associations in shaping infra-communities.

Our capacity to predict the moose gut microbial community at the population scale increased when we included spatial information, indicating that either stochastic events such as microbial dispersal or spatial similarities in diet may be important in structuring associations between gut microbes. Dispersal of microbes is thought to be passive (Nemergut et al., 2013), however, these microbial species are often in high abundance and have broad distributions making dispersal challenging to quantify (Evans et al., 2017; Zhou & Ning, 2017). Biogeographical studies have shown that dispersal limitation is important for structuring microbial communities including those of the gut (Evans et al., 2017; Hanson, Fuhrman, Horner-Devine, & Martiny, 2012; Moeller et al., 2017), but rarely over a relatively small area within a host population. What role moose diet explicitly played in shaping our moose gut microbial communities is still an open question. Future studies linking host microbial data to measures of host diet, such as stable isotope analysis (Hofman-Kamińska, Bocherens, Borowik, Drucker, & Kowalczyk, 2018), will enable dispersal limitation and diet to be decoupled in future models. Nonetheless, we show that including host spatial data in future work is necessary for robustly quantifying microbial co-occurrence patterns.

We also found strong positive and negative associations between OTUs from different phyla and divergent functional groups in the moose gut microbial community. Clostridia (Firmicutes) and Mollicutes (Tenericutes) were network ‘hubs’ involved in a relatively high number of positive associations in our moose population, and often with each other (Table 1). Clostridia OTUs are also hubs for associations in the human gut community (Banerjee, Schlaeppi, & van der Heijden, 2018; Faust et al., 2012), but the importance of Mollicutes has not been, to our knowledge, reported elsewhere. Moreover, we detected no negative associations between Bacteroidetes and Firmicutes even though Bacteroidetes was common in our samples; this group may not play a dominant role in structuring moose gut microbial communities as they do in humans (Banerjee et al., 2018; Faust et al., 2012). Whether the same relationships apply in herbivore populations where Bacteroidetes is dominant, as was the case in an Alaskan and Swedish moose (Ishaq & Wright, 2014; Svartström et al., 2017), remains an open question. More generally, Faust et al. (2012) found an increased likelihood of negative associations between OTUs that were functionally and phylogenetically dissimilar (with opposite true for more related OTUs). We did not see such a pattern in our co-occurrence network and why we would get such a different result in moose is unclear. The Faust dataset consisted of 240 individuals from the Human Microbiome Project (Methé et al., 2012), but large methodological differences make direct comparison difficult, as factors such as spatial relationships between subjects were not quantified. How robust these relationships are also likely to be impacted by taxonomic resolution of each network (Faust et al., 2012). As we used 16S rRNA gene sequencing, we were unable to get sequence identifications to species level. Further, functional profiles were based on predictions from reference genomes, not direct identification of functional genes or proteins (Langille et al., 2013). Shotgun metagenomic sequencing could allow for both classification sequences to species-level and provide much more detailed insights into the functional patterns shaping microbial species associations.

Here we have demonstrated that graphic network models can untangle how interspecific interactions can shape gut microbial communities in moose. Additionally, we show that MRF and CRF models can robustly construct co-occurrence networks (Clark, Wells, & Lindberg, 2018b) and, coupled with taxonomic and functional information, find high-resolution insights into interspecific associations. Graphical network models, as with other correlation-based approaches, do have limitations. Correlations identified by techniques such as MRF or CRF should be treated with caution as they do not imply causation (e.g., Barner et al., 2018; Dormann et al., 2018). Follow up analysis with tools such as structural equation modelling (SEM) could be used to go beyond correlations to explore potential causal relationships between microbes (Banerjee et al., 2018). Nonetheless, by explicitly incorporating spatial data and covariates into graphical models, our method offers a step forward in characterising associations between species, and we envisage this method will be broadly useful for researchers working on micro- and macro-community dynamics alike. Studies analysing associations between infra-community microbial species are rare in wildlife, even though they can provide important insights into the ecological dynamics operating within or on the host. Given the decreasing cost of microbial surveys and analytical advances such as ours, studies that can disentangle microbial infra-community dynamics in wildlife species will become more frequent, and this can ultimately provide a more nuanced understanding of wildlife health.

## Acknowledgements

This material is based upon work supported by the Cooperative State Research Service, U.S. Department of Agriculture, under Project No. MINV-62-051.

## Data accessibility

All data will be made accessible in Dryad.

## Author contribution statement

NFJ, MC, TW, JF, TJ, AK and MEC came up with the project design. NFJ, NC and EM conducted the analysis. NC coded the spatial MRF/CRF functions. MC, TW, SM and JF provided data. All authors contributed critically to the drafts and gave final approval for publication.

## References

Antwis, R. E., Griffiths, S. M., Harrison, X. A., Aranega-Bou, P., Arce, A., Bettridge, A. S., … Sutherland, W. J. (2017). Fifty important research questions in microbial ecology. FEMS Microbiology Ecology, 93(5). doi:10.1093/femsec/fix044

Aragón, P., Carrascal, L. M., & Palomino, D. (2018). Macro-spatial structure of biotic interactions in the distribution of a raptor species. Journal of Biogeography, 45(8), 1859–1871. doi:10.1111/jbi.13389

Araújo, M. B., & Rozenfeld, A. (2014). The geographic scaling of biotic interactions. Ecography, 37(5), 406–415. doi:10.1111/j.1600-0587.2013.00643.x

Banerjee, S., Schlaeppi, K., & van der Heijden, M. G. A. (2018). Keystone taxa as drivers of microbiome structure and functioning. Nature Reviews Microbiology, 16(9), 567–576. doi:10.1038/s41579-018-0024-1

Barner, A. K., Coblentz, K. E., Hacker, S. D., & Menge, B. A. (2018). Fundamental contradictions among observational and experimental estimates of non-trophic species interactions. Ecology, 99(3), 557–566. doi:10.1002/ecy.2133

Bauer, E., & Thiele, I. (2018). From network analysis to functional metabolic modeling of the human gut microbiota. MSystems, 3(3), e00209-17. doi:10.1128/mSystems.00209-17

Bolger, A. M., Lohse, M., & Usadel, B. (2014). Trimmomatic: a flexible trimmer for Illumina sequence data. Bioinformatics, 30(15), 2114–2120. doi:10.1093/bioinformatics/btu170

Britton, R. A., & Young, V. B. (2014). Role of the intestinal microbiota in resistance to colonization by Clostridium difficile. Gastroenterology, 146(6), 1547–1553. doi:10.1053/j.gastro.2014.01.059

Butler, E., Carstensen, M., Hildebrand, E., & Giudice, J. (2012). Northeast Minnesota moose herd health assessment 2007–2012. In L. Cornicelli, M. Carstensen, M. Grund, M. Larson, & Lawrence JS (Eds.), Summaries of wildlife research findings 2012. St Paul: Minnesota Department of Natural Resources. Retrieved from http://www.mndnr.gov

Camp, J. G., Kanther, M., Semova, I., & Rawls, J. F. (2009). Patterns and scales in gastrointestinal microbial ecology. Gastroenterology, 136(6), 1989–2002. doi:10.1053/j.gastro.2009.02.075

Caporaso, J. G., Bittinger, K., Bushman, F. D., DeSantis, T. Z., Andersen, G. L., & Knight, R. (2010). PyNAST: a flexible tool for aligning sequences to a template alignment. Bioinformatics, 26(2), 266–267. doi:10.1093/bioinformatics/btp636

Caporaso, J. G., Kuczynski, J., Stombaugh, J., Bittinger, K., Bushman, F. D., Costello, E. K., … Knight, R. (2010). QIIME allows analysis of high-throughput community sequencing data. Nature Methods, 7(5), 335–336. doi:10.1038/nmeth.f.303

Carstensen, M., Hildebrand, E. C., Plattner, D., Dexter, M., St-Louis, V., Jennelle, C., & Wright, R. G. (2018). Determining cause specific mortality of adult moose in Northeast Minnesota, February 2013 - July 2017. Retrieved from https://files.dnr.state.mn.us/wildlife/research/summaries/health/2016_moosemortality.pdf

Chase, J. M., & Myers, J. A. (2011). Disentangling the importance of ecological niches from stochastic processes across scales. Philosophical Transactions of the Royal Society B: Biological Sciences, 366(1576), 2351–2363. doi:10.1098/rstb.2011.0063

Cheng, Y., Fox, S., Pemberton, D., Hogg, C., Papenfuss, A. T., & Belov, K. (2015). The Tasmanian devil microbiome—implications for conservation and management. Microbiome, 3(1), 76. doi:10.1186/s40168-015-0143-0

Clark, N. J., Wells, K., & Lindberg, O. (2018). MRFcov: Markov Random Fields with additional covariates. R package version 1.0.33. Availabe at GitHub. https://github.com/nicholasjclark/MRFcov.

Clark, N. J., Wells, K., & Lindberg, O. (2018). Unravelling changing interspecific interactions across environmental gradients using Markov random fields. Ecology, 99(6), 1277–1283. doi:10.1002/ecy.2221

Csárdi, G., & Nepusz, T. (2006). The igraph software package for complex network research. InterJournal Complex Systems, 1695.

de Araújo, C. B., Marcondes-Machado, L. O., & Costa, G. C. (2014). The importance of biotic interactions in species distribution models: a test of the Eltonian noise hypothesis using parrots. Journal of Biogeography, 41(3), 513–523. doi:10.1111/jbi.12234

Delgiudice, G. D. (2018). 2018 Aerial Moose Survey. Retrieved from https://files.dnr.state.mn.us/wildlife/moose/moosesurvey.pdf

DeSantis, T. Z., Hugenholtz, P., Larsen, N., Rojas, M., Brodie, E. L., Keller, K., … Andersen, G. L. (2006a). Greengenes, a chimera-checked 16S rRNA gene database and workbench compatible with ARB. Applied and Environmental Microbiology, 72(7), 5069–72. doi:10.1128/AEM.03006-05

DeSantis, T. Z., Hugenholtz, P., Larsen, N., Rojas, M., Brodie, E. L., Keller, K., … Andersen, G. L. (2006b). Greengenes, a chimera-checked 16S rRNA gene database and workbench compatible with ARB. Applied and Environmental Microbiology, 72(7), 5069–5072. doi:10.1128/AEM.03006-05

Dormann, C. F., Bobrowski, M., Dehling, D. M., Harris, D. J., Hartig, F., Lischke, H., … Kraan, C. (2018). Biotic interactions in species distribution modelling: 10 questions to guide interpretation and avoid false conclusions. Global Ecology and Biogeography. doi:10.1111/geb.12759

Edgar, R. C. (2010). Search and clustering orders of magnitude faster than BLAST. Bioinformatics, 26(19), 2460–2461. doi:10.1093/bioinformatics/btq461

Evans, S., Martiny, J. B. H., & Allison, S. D. (2017). Effects of dispersal and selection on stochastic assembly in microbial communities. The ISME Journal, 11(1), 176–185. doi:10.1038/ismej.2016.96

Faust, K., Sathirapongsasuti, J. F., Izard, J., Segata, N., Gevers, D., Raes, J., & Huttenhower, C. (2012). Microbial co-occurrence relationships in the human microbiome. PLoS Computational Biology, 8(7), e1002606. doi:10.1371/journal.pcbi.1002606

Ganz, H. H., Doroud, L., Firl, A. J., Hird, S. M., Eisen, J. A., & Boyce, W. M. (2017). Community-level differences in the microbiome of healthy wild mallards and those infected by Influenza A riruses. MSystems, 2(1), e00188-16. doi:10.1128/mSystems.00188-16

Gohl, D. M., Vangay, P., Garbe, J., MacLean, A., Hauge, A., Becker, A., … Beckman, K. B. (2016). Systematic improvement of amplicon marker gene methods for increased accuracy in microbiome studies. Nature Biotechnology, 34(9), 942–949. doi:10.1038/nbt.3601

Hanson, C. A., Fuhrman, J. A., Horner-Devine, M. C., & Martiny, J. B. H. (2012). Beyond biogeographic patterns: processes shaping the microbial landscape. Nature Reviews Microbiology, 10(7), 497–506. doi:10.1038/nrmicro2795

Harris, D. J. (2016). Inferring species interactions from co-occurrence data with Markov networks. Ecology, 97(12), 3308–3314. doi:10.1002/ecy.1605

Henderson, G., Cox, F., Ganesh, S., Jonker, A., Young, W., Global Rumen Census Collaborators, G. R. C., & Janssen, P. H. (2015). Rumen microbial community composition varies with diet and host, but a core microbiome is found across a wide geographical range. Scientific Reports, 5, 14567. doi:10.1038/srep14567

Herren, C. M., & McMahon, K. D. (2018). Keystone taxa predict compositional change in microbial communities. Environmental Microbiology, 20(6), 2207–2217. doi:10.1111/1462-2920.14257

Hofman-Kamińska, E., Bocherens, H., Borowik, T., Drucker, D. G., & Kowalczyk, R. (2018). Stable isotope signatures of large herbivore foraging habitats across Europe. PLOS ONE, 13(1), e0190723. doi:10.1371/journal.pone.0190723

Hooper, L. V., Littman, D. R., & Macpherson, A. J. (2012). Interactions between the microbiota and the immune system. Science, 336(6086), 1268–1273. doi:10.1126/science.1223490

Hundertmark, K. J., Bowyer, R. T., Shields, G. F., & Schwartz, C. C. (2003). Mitochondrial phylogeography of moose (Alces alces) in North America. Journal of Mammalogy, 84(2), 718–728. doi:10.1644/1545-1542(2003)084<0718:MPOMAA>2.0.CO;2

Ishaq, S. L., Sundset, M. A., Crouse, J., & Wright, A.-D. G. (2015). High-throughput DNA sequencing of the moose rumen from different geographical locations reveals a core ruminal methanogenic archaeal diversity and a differential ciliate protozoal diversity. Microbial Genomics, 1(4), e000034. doi:10.1099/mgen.0.000034

Ishaq, S. L., & Wright, A.-D. G. (2012). Insight into the bacterial gut microbiome of the North American moose (Alces alces). BMC Microbiology, 12(1), 212. doi:10.1186/1471-2180-12-212

Ishaq, S. L., & Wright, A. D. (2014). High-throughput DNA sequencing of the ruminal bacteria from moose (Alces alces) in Vermont, Alaska, and Norway. Microbial Ecology, 68(2), 185–195. doi:10.1007/s00248-014-0399-0

Jani, A. J., & Briggs, C. J. (2014). The pathogen Batrachochytrium dendrobatidis disturbs the frog skin microbiome during a natural epidemic and experimental infection. Proceedings of the National Academy of Sciences, 111(47), E5049–E5058. doi:10.1073/pnas.1412752111

Jani, A. J., & Briggs, C. J. (2018). Host and aquatic environment shape the amphibian skin microbiome but effects on downstream resistance to the pathogen Batrachochytrium dendrobatidis are variable. Frontiers in Microbiology, 9(MAR), 487. doi:10.3389/fmicb.2018.00487

Kammann, E. E., & Wand, M. P. (2003). Geoadditive models. Journal of the Royal Statistical Society: Series C (Applied Statistics), 52(1), 1–18. doi:10.1111/1467-9876.00385

Kanehisa, M., Sato, Y., Kawashima, M., Furumichi, M., & Tanabe, M. (2016). KEGG as a reference resource for gene and protein annotation. Nucleic Acids Research, 44(D1), D457–D462. doi:10.1093/nar/gkv1070

Kohl, K. D., Varner, J., Wilkening, J. L., & Dearing, M. D. (2018). Gut microbial communities of American pikas (Ochotona princeps): Evidence for phylosymbiosis and adaptations to novel diets. Journal of Animal Ecology, 87(2), 323–330. doi:10.1111/1365-2656.12692

Langille, M. G. I., Zaneveld, J., Caporaso, J. G., McDonald, D., Knights, D., Reyes, J. A., … Huttenhower, C. (2013). Predictive functional profiling of microbial communities using 16S rRNA marker gene sequences. Nature Biotechnology, 31(9), 814–821. doi:10.1038/nbt.2676

Le Borgne, H., Hébert, C., Dupuch, A., Bichet, O., Pinaud, D., & Fortin, D. (2018). Temporal dynamics in animal community assembly during post-logging succession in boreal forest. PLOS ONE, 13(9), e0204445. doi:10.1371/journal.pone.0204445

Lee, J. D., & Hastie, T. J. (2015). Learning the structure of mixed graphical models. Journal of Computational and Graphical Statistics, 24(1), 230–253. doi:10.1080/10618600.2014.900500

Lenarz, M. S. (2009). 2009 Aerial Moose Survey. St Paul, USA.

Ley, R. E., Hamady, M., Lozupone, C., Turnbaugh, P. J., Ramey, R. R., Bircher, J. S., … Gordon, J. I. (2008). Evolution of mammals and their gut microbes. Science, 320(5883), 1647–1651. doi:10.1126/science.1155725

Masella, A. P., Bartram, A. K., Truszkowski, J. M., Brown, D. G., & Neufeld, J. D. (2012). PANDAseq: paired-end assembler for illumina sequences. BMC Bioinformatics, 13(1), 31. doi:10.1186/1471-2105-13-31

McKenney, E. A., Williamson, L., Yoder, A. D., Rawls, J. F., Bilbo, S. D., & Parker, W. (2015). Alteration of the rat cecal microbiome during colonization with the helminth Hymenolepis diminuta. Gut Microbes, 6(3), 182–193. doi:10.1080/19490976.2015.1047128

Methé, B. A., Nelson, K. E., Pop, M., Creasy, H. H., Giglio, M. G., Huttenhower, C., … White, O. (2012). A framework for human microbiome research. Nature, 486(7402), 215–221. doi:10.1038/nature11209

Moeller, A. H., Peeters, M., Ayouba, A., Ngole, E. M., Esteban, A., Hahn, B. H., & Ochman, H. (2015). Stability of the gorilla microbiome despite simian immunodeficiency virus infection. Molecular Ecology, 24(3), 690–697. doi:10.1111/mec.13057

Moeller, A. H., Suzuki, T. A., Lin, D., Lacey, E. A., Wasser, S. K., & Nachman, M. W. (2017). Dispersal limitation promotes the diversification of the mammalian gut microbiota. Proceedings of the National Academy of Sciences of the United States of America, 114(52), 13768–13773. doi:10.1073/pnas.1700122114

Mshelia, E. S., Adamu, L., Wakil, Y., Turaki, U. A., Gulani, I. A., & Musa, J. (2018). The association between gut microbiome, sex, age and body condition scores of horses in Maiduguri and its environs. Microbial Pathogenesis, 118, 81–86. doi:10.1016/j.micpath.2018.03.018

Muegge, B. D., Kuczynski, J., Knights, D., Clemente, J. C., González, A., Fontana, L., … Gordon, J. I. (2011). Diet drives convergence in gut microbiome functions across mammalian phylogeny and within humans. Science, 332(6032), 970–4. doi:10.1126/science.1198719

Murray, D. L., Cox, E. W., Ballard, W. B., Whitlaw, H. A., Lenarz, M. S., Custer, T. W., … Fuller, T. K. (2006). Pathogens, nutritional deficiency, and climate influences on a declining moose population. Wildlife Monographs, 166(1), 1–30. doi:10.2193/0084-0173(2006)166[1:PNDACI]2.0.CO;2

Näpflin, K., & Schmid-Hempel, P. (2018). Host effects on microbiota community assembly. Journal of Animal Ecology, 87(2), 331–340. doi:10.1111/1365-2656.12768

Nemergut, D. R., Schmidt, S. K., Fukami, T., O’Neill, S. P., Bilinski, T. M., Stanish, L. F., … Ferrenberg, S. (2013). Patterns and processes of microbial community assembly. Microbiology and Molecular Biology Reviews: MMBR, 77(3), 342–56. doi:10.1128/MMBR.00051-12

Oksanen, J., Blanchet, F. G., Kindt, R., Legendre, P., Minchin, P. R., O’Hara, R. B., … Wagner, H. (2013). Package ‘vegan.’ R Package Ver. 2.0–8, 254.

Ovaskainen, O., Abrego, N., Halme, P., & Dunson, D. (2016). Using latent variable models to identify large networks of species-to-species associations at different spatial scales. Methods in Ecology and Evolution, 7(5), 549–555. doi:10.1111/2041-210X.12501

Ovaskainen, O., Tikhonov, G., Norberg, A., Guillaume Blanchet, F., Duan, L., Dunson, D., … Abrego, N. (2017). How to make more out of community data? A conceptual framework and its implementation as models and software. Ecology Letters, 20(5), 561–576. doi:10.1111/ele.12757

Severud, W. (2017). Assessing calf survival and the quantitative impact of reproductive success on the declining moose (Alces alces) population in northeastern Minnesota. University of Minnesota. Retrieved from https://conservancy.umn.edu/handle/11299/191446

Svartström, O., Alneberg, J., Terrapon, N., Lombard, V., De Bruijn, I., Malmsten, J., … Andersson, A. F. (2017). Ninety-nine de novo assembled genomes from the moose (Alces alces) rumen microbiome provide new insights into microbial plant biomass degradation. ISME Journal, 11(11), 2538–2551. doi:10.1038/ismej.2017.108

Vellend, M., Srivastava, D. S., Anderson, K. M., Brown, C. D., Jankowski, J. E., Kleynhans, E. J., … Xue, X. (2014). Assessing the relative importance of neutral stochasticity in ecological communities. Oikos, 123(12), 1420–1430. doi:10.1111/oik.01493

Wood, S. N. (2003). Thin plate regression splines. Journal of the Royal Statistical Society: Series B (Statistical Methodology), 65(1), 95–114. doi:10.1111/1467-9868.00374

Wünschmann, A., Armien, A. G., Butler, E., Schrage, M., Stromberg, B., Bender, J. B., … Carstensen, M. (2015). Necropsy findings in 62 opportunistically collected free-ranging moose (Alces alces) from Minnesota, USA (2003–13). Journal of Wildlife Diseases, 51(1), 157–165. doi:10.7589/2014-02-037

Zelezniak, A., Andrejev, S., Ponomarova, O., Mende, D. R., Bork, P., & Patil, K. R. (2015). Metabolic dependencies drive species co-occurrence in diverse microbial communities. Proceedings of the National Academy of Sciences of the United States of America, 112(20), 6449–54. doi:10.1073/pnas.1421834112

Zhou, J., & Ning, D. (2017). Stochastic community assembly: Does it matter in Microbial Ecology? Microbiology and Molecular Biology Reviews: MMBR, 81(4), e00002-17. doi:10.1128/MMBR.00002-17

