## Supplemental figures and tables for "Microbial associations and spatial proximity predict North American moose (*Alces alces*) gastrointestinal community composition"

**Supplementary materials**

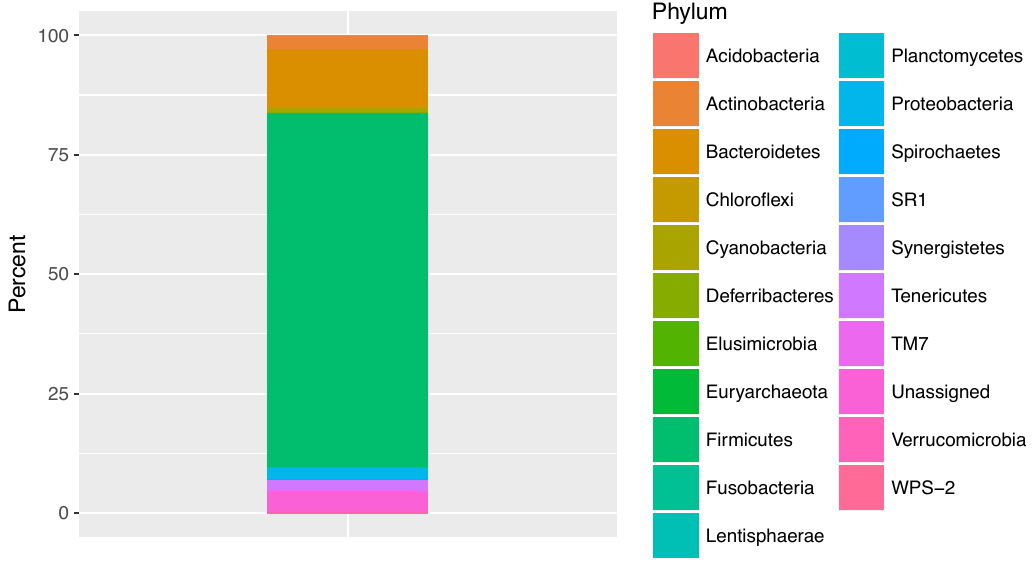

**Figure S1:** Relative abundance of OTUs at the phylum level for all samples (n = 55). Common and rare OTUs were included in this figure.

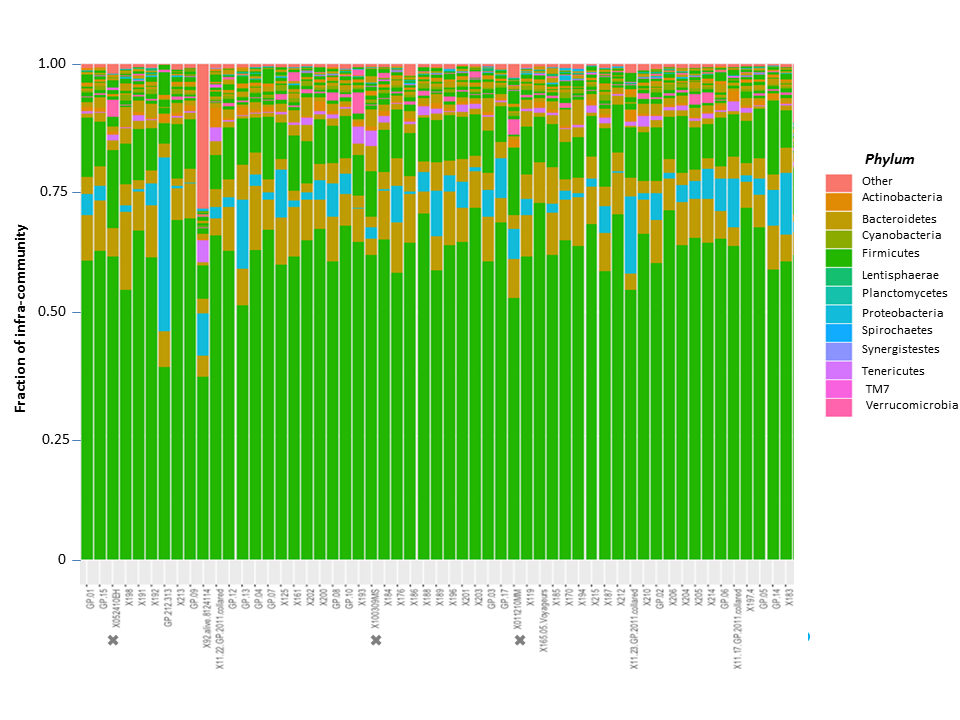

**Figure S2:** Proportion of microbial phyla sampled from each moose. Crosses indicate that these individuals were not considered in the analyses presented in the main text. Common and rare OTUs were included in this figure.

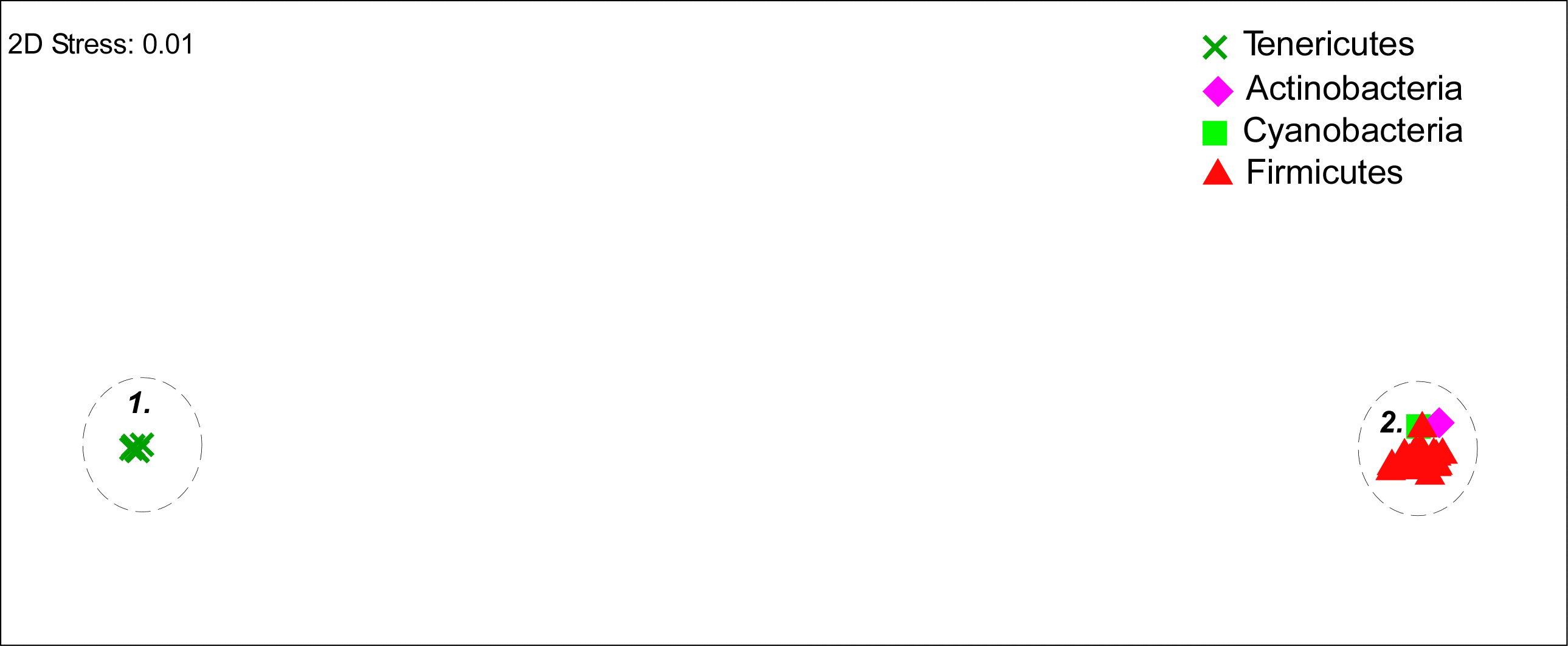

**Figure S3:** Non-metric multidimensional relationships showing the functional relationships between the remaining (i.e., non-filtered) OTUs in multidimensional space. Different symbols refer to different taxonomic groups (see key). Dashed circles indicate distinct functional groups (FGs), with FG1 and FG2 illustrated here.

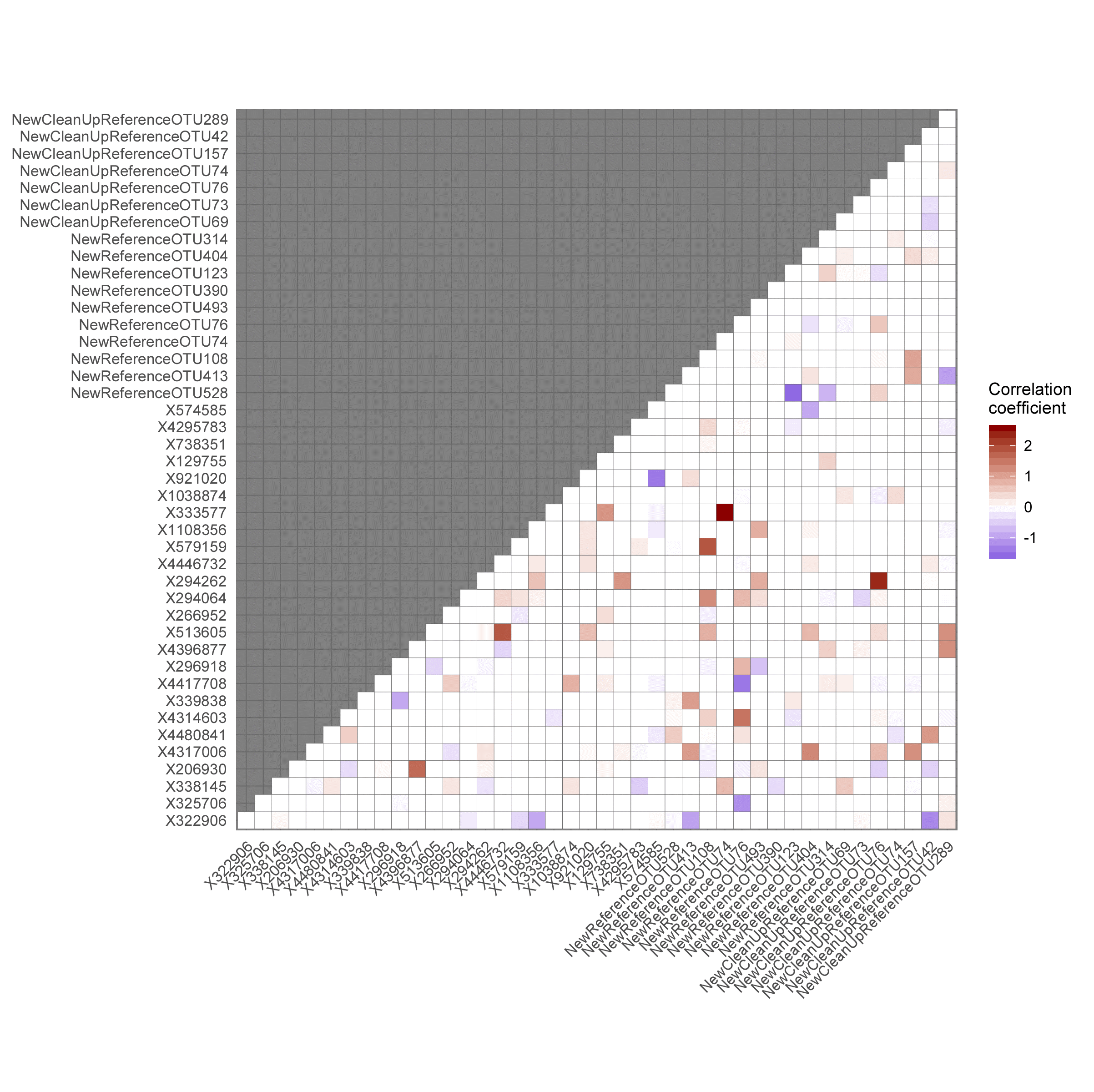

**Figure S4:** Co-occurrence matrix displaying the estimated co-occurrence coefficients between each pair of OTUs.
