## Appendix S1 for "Microbial associations and spatial proximity predict North American moose (*Alces alces*) gastrointestinal community composition"

### Microbiome Analysis

Read in the BIOM file from QIIME and convert to a matrix

```
library(biomformat)
library(OTUtable)
library(gradientForest)
library(missForest)
dat <- read_biom("Data/otu_table_rarefied.biom")
otu_table <- as.data.frame(as.matrix(biom_data(dat)))
```

Gather metadata

```
taxonomy <- observation_metadata(dat)
metadata <- sample_metadata(dat)
```

Remove duplicates at the same level. Level options are from column names from taxonomic dataset. No need to do this

```
combine_otus("Genus", table, taxonomy)
write.csv(otu_table, file = "Data/otuDataTable.csv")
```

To make a table containing only phyla with at least 10% abundance in any one sample and were observed at any abundance in at least 10% of samples. Convert the table to presence / absence and remove OTUs that are too common for analysis

```
otuFilter <- filter_taxa(otu_table, abundance = 0.01,
    persistence = 5)
otuFilter[otuFilter > 0] <- 1
otuFilter$new <- rowSums(otuFilter[, 1:55])
otuFilterNoCommon <- subset(otuFilter, new <
    45)
write.csv(otuFilterNoCommon, file = "Data/OTUtableFiltered.csv")
```

Here, we use MDS to focus on OTUs that were driving the compostional patterns. MDS Scores with `p > 0.45` Pearson correlation coefficents are filtered out

```
dat <- read.csv("Data/OTUtableFiltered.csv",
    head = T, row.names = 1)
mds <- read.csv("Data/MDSscores.csv", head = T)
a <- names(mds)
OTUReduced <- dat[which(names(dat) %in% a)]
OTUReduced <- OTUReduced[order(row.names(OTUReduced)),
    ]
OTUReduced <- OTUReduced[-c(1:3), ]
OTUReduced <- OTUReduced[, -41]
```

Import environmental covariate data, merge with the OTU data and save

```
env <- read.csv("Data/envAll.csv", head = T,
    row.names = 1)
env$Sex <- as.factor(env$Sex)
env <- env[order(row.names(env)), ]
env$Sex <- as.integer(env$Sex)
mooseMB <- cbind(OTUReduced, env)
write.csv(mooseMB, file = "Data/mooseMB.csv")
```

Load `MRFcov` for running Conditional Random Fields co-occurrence analysis. For this model, very rare OTUs (occurring in fewer than 5% of observations) will need to be removed

```
library(MRFcov)
mooseMB$New.CleanUp.ReferenceOTU82 <- NULL
```

Extract coordinates columns to use for incorporating spatial splines as covariates in the model (for now, these need to be named `Latitude` and `Longitude`)

```
Latitude <- mooseMB$Capture.Location.UTM.Easting
mooseMB$Capture.Location.UTM.Easting <- NULL
Longitude <- mooseMB$Capture.Location.UTM.Northing
mooseMB$Capture.Location.UTM.Northing <- NULL
coords <- (data.frame(Latitude = Latitude,
    Longitude = Longitude))
```

Prep the remaining covariates for CRF analysis. Here, we change categorical covariates to model matrix format and scale numeric covariates to unit variance

```
mooseMB$Sex <- as.factor(mooseMB$Sex)

library(dplyr)
analysis.data <- mooseMB %>% cbind(., data.frame(model.matrix(~.[,
    "Sex"], .)[, -1])) %>% dplyr::select(-Sex) %>%
    dplyr::rename_all(funs(gsub("\\.|model.matrix",
        "", .))) %>% dplyr::mutate_at(vars(CaptureDate:BodyCondition),
    funs(as.vector(scale(.))))
```

We also create a dataset for non-spatial analysis for comparisons. For this, we add the `Latitude` and `Longitude` columns back in and scale them to unit variance

```
analysis.data.nonspatial <- analysis.data %>%
    dplyr::bind_cols(data.frame(Latitude = Latitude,
        Longitude = Longitude)) %>% dplyr::mutate_at(vars(Latitude,
    Longitude), funs(as.vector(scale(.))))
```

Now we generate MRF and CRF models to determine which model fits the data most appropriately. First, the nonspatial MRF (without covariates)

```
moose.mrf <- MRFcov(data = analysis.data.nonspatial[,
    1:42], n_nodes = 42, family = "binomial",
    n_cores = 3)
```

Use this model to generate predictions

```
mrf.predictions <- predict_MRF(data = analysis.data.nonspatial[,
    1:42], MRF_mod = moose.mrf)
```

Test how well the MRF predicts the data by comparing the predicted to the observed values. Here, we split the data into five folds and test for model specificity, sensitivity, positive predictive value and proportion of true predictions using the `cv_MRF_diag` function

```
mrf.cv <- lapply(seq_len(100), function(x) {
    cv_MRF_diag(data = analysis.data.nonspatial[,
        1:42], n_nodes = 42, n_folds = 5,
        n_cores = 1, family = "binomial",
        compare_null = FALSE, plot = FALSE,
        cached_model = moose.crf, cached_predictions = list(predictions = mrf.predictions),
        sample_seed = 10)
})
mrf.cv <- do.call(rbind, mrf.cv)
```

Next, we repeat the above for a nonspatial CRF (including environmental covariates, but without spatial splines)

```
moose.crf <- MRFcov(data = analysis.data.nonspatial,
    n_nodes = 42, family = "binomial", n_cores = 3)
crf.predictions <- predict_MRF(data = analysis.data.nonspatial[,
    1:42], MRF_mod = moose.crf)
crf.cv <- lapply(seq_len(100), function(x) {
    cv_MRF_diag(data = analysis.data.nonspatial,
        n_nodes = 42, n_folds = 5, n_cores = 1,
        family = "binomial", compare_null = FALSE,
        plot = FALSE, cached_model = moose.crf,
        cached_predictions = list(predictions = crf.predictions),
        sample_seed = 10)
})
crf.cv <- do.call(rbind, crf.cv)
```

Next, we fit the spatial CRF. Here, the coordinates are used to produce spatial regression splines with the call `mgcv::smooth.construct2(object = mgcv::s(Latitude, Longitude, bs = "gp"), data = coords, knots = NULL)`. Splines are a series of different equations pieced together, which tends to increase the accuracy of interpolated values at the cost of the ability to project outside the data range. These are highly appropriate here, as we are not interested in using them for prediction but more to account for non-independence in our observations.

```
moose.crf.spatial <- MRFcov_spatial(data = analysis.data,
    n_nodes = 42, family = "binomial", coords = coords,
    n_cores = 3)
spatial.predictions <- predict_MRF(data = moose.crf.spatial$mrf_data,
    prep_covariates = F, MRF_mod = moose.crf.spatial)
spatial.cv <- lapply(seq_len(100), function(x) {
    cv_MRF_diag(data = analysis.data, n_nodes = 42,
        n_folds = 5, n_cores = 1, family = "binomial",
        compare_null = FALSE, plot = FALSE,
        cached_model = moose.crf.spatial,
        cached_predictions = list(predictions = spatial.predictions),
        sample_seed = 10)
})
spatial.cv <- do.call(rbind, spatial.cv)
```

Now that we have run all of the models and generated predictions, we can examine the proportion of unique observations that were correctly predicted by each of the different models

```
quantile(mrf.cv$mean_tot_pred, probs = c(0.025,
    0.5, 0.975))
```

```
##      2.5%       50%     97.5%
## 0.8035714 0.8280423 0.8503401
```

```
quantile(crf.cv$mean_tot_pred, probs = c(0.025,
    0.5, 0.975))
```

```
##      2.5%       50%     97.5%
## 0.7131494 0.7489177 0.7809524
```

```
quantile(spatial.cv$mean_tot_pred, probs = c(0.025,
    0.5, 0.975))
```

```
##      2.5%       50%     97.5%
## 0.7595238 0.7876221 0.8165051
```

Interestingly, it seems that a standard MRF (without covariates) fits the data better than either of the other models (though inclusion of the spatial term clearly improves the fit of the CRF), suggesting that biotic interactions are much more important than environmental or spatial covariates for predicting OTU occurrence probabilities. This could also be reflecting that our 52 observations are not adequate to detect changing interactions across environmental gradients, though it is difficult to know until we can acquire more data.

While the above results suggest the MRF is a better fit, simply focussing on proportions of true predictions can be misleading, as it could be possible that one model does a good job of predicting absences while another can predict presences better. To differentiate, we can explore predictive capacities in more depth by plotting a range of predictive metrics for the MRF and CRF models

```
library(gridExtra)
cv_MRF_diag_rep(data = analysis.data.nonspatial,
    n_nodes = 42, n_cores = 3, family = "binomial",
    compare_null = T, plot = T, n_fold_runs = 100)
```

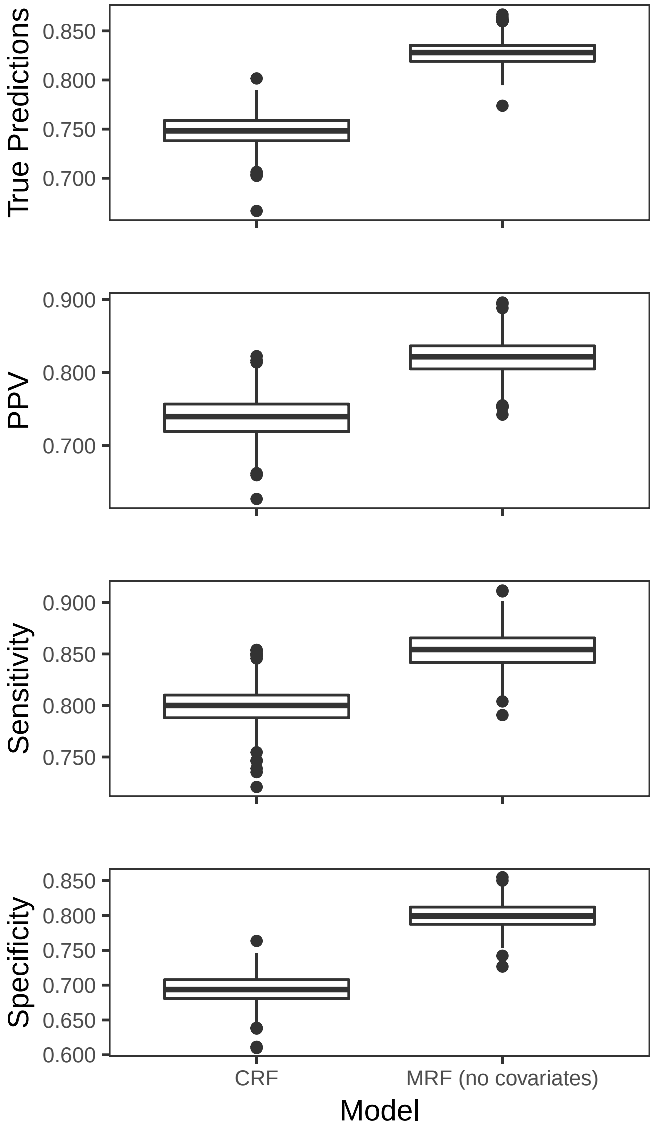

All of these metrics suggest that the MRF is the better model for this dataset and that addition of environmental covariates does not improve fit, which is encouraging as this means we do not need to rely on more complex models to understand co-occurrence patterns. A logical next step here is to test if adding the spatial splines to the MRF can improve fit even further, as it may still make sense to account for spatial processes

```
cv_MRF_diag_rep_spatial(data = analysis.data[,
    1:42], n_nodes = 42, family = "binomial",
    coords = coords, n_cores = 3, compare_null = T,
    plot = T, n_fold_runs = 100)
```

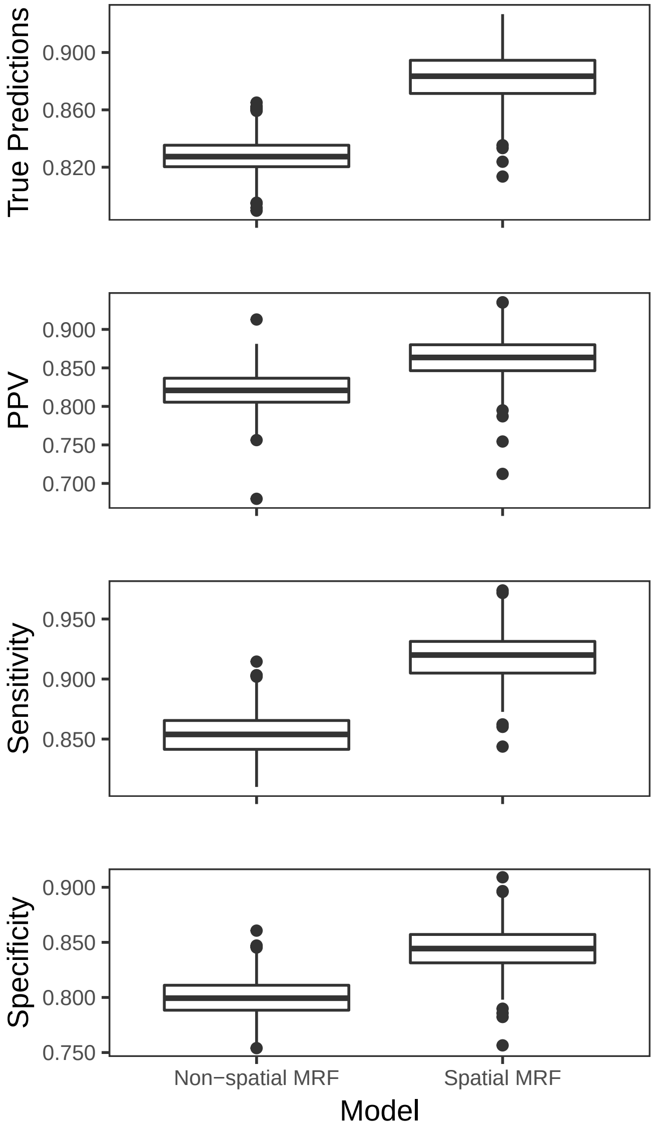


This results suggest that the spatial splines further improve the model's predictive capacity, so we can stick to this model for interpretation.

We can now inspect the spatial MRF's regression coefficients to identify possible biotic interactions, here plotted as a heatmap

```
moose.mrf.spatial <- MRFcov_spatial(data = analysis.data[,
    1:42], n_nodes = 42, family = "binomial",
    coords = coords, n_cores = 3)
```

```
plotMRF_hm(moose.mrf.spatial, main = "Estimated OTU Co-occurrence Probabilities")
```

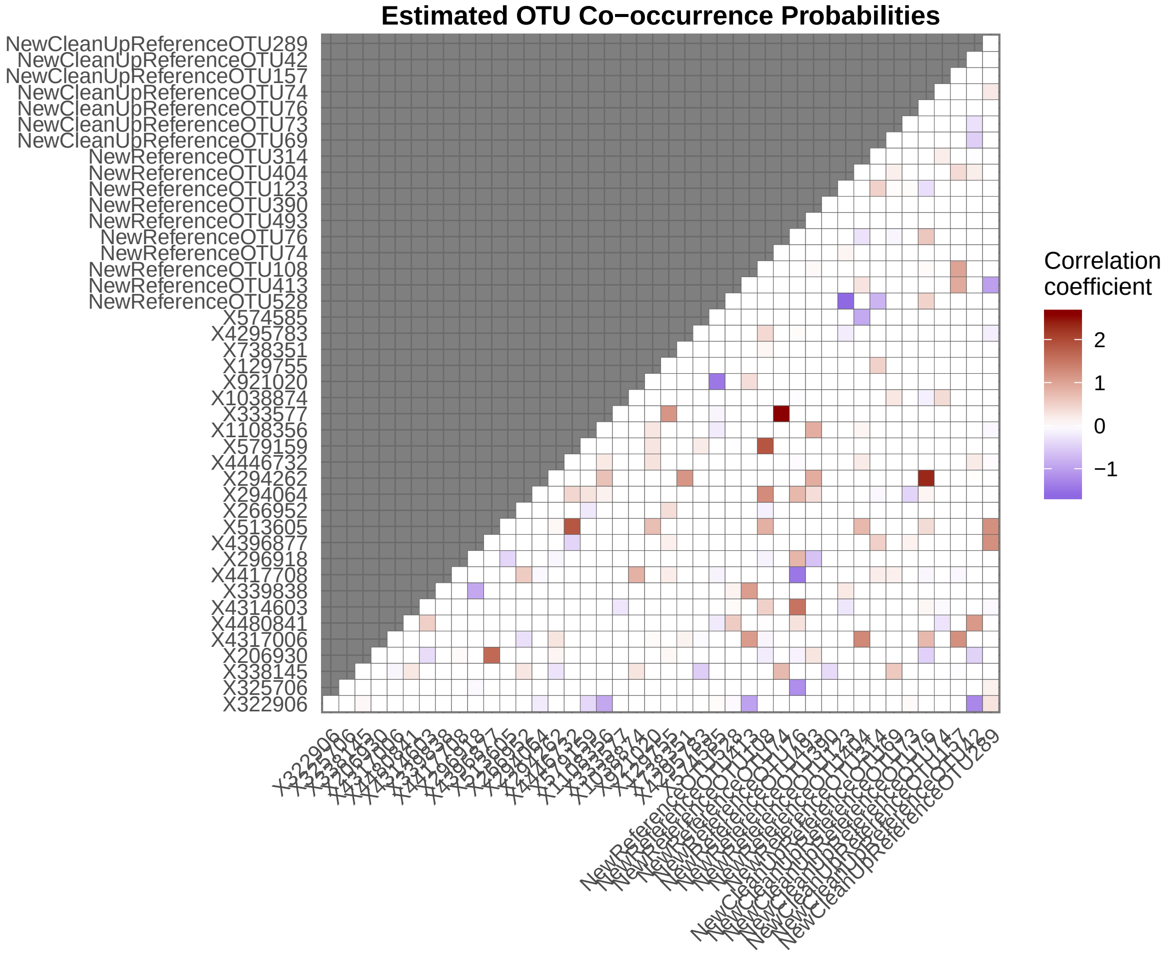

It is worth bootstrapping the spatial MRF to capture uncertainty in model parameters. This function will randomly shuffle observations and fit the spatial MRF in each bootstrap iteration. We perform 100 iterations to calculate the spread of model coefficients

```
moose.bootstrap.spatial <- bootstrap_MRF(data = analysis.data[,
    1:42], n_nodes = 42, family = "binomial",
    spatial = TRUE, coords = coords, n_cores = 3,
    n_bootstraps = 100)
```

Finally, we can calculate average occurrence probability for each OTU. Because we use logistic regression with a logit link function to estimate occurrences in the MRF, we can take the inverse logit of intercepts to represent the 'average' occurrence probability of each OTU. We also add PiCRUST taxonomy information and summarise the numbers of positive and negative interactions that were identified for each OTU to gain insight into their respective influence on microbiome composition

```
taxonomy <- readxl::read_xlsx("Data/otu_taxonomy.xlsx",
    sheet = 1) %>% dplyr::mutate(Group = paste(phylum,
    class, sep = ":")) %>% dplyr::mutate(OTU = gsub("[.]",
    "", OTU_IDs)) %>% dplyr::select(OTU,
    Group)

summary_table <- data.frame(OTU = rownames(moose.mrf.spatial$graph),
    Ave.occurrence = round(boot::inv.logit(moose.bootstrap.spatial$direct_coef_means[,
        1]), 4), Pos.interactions = apply(moose.bootstrap.spatial$direct_coef_means[1:42,
        2:43], 1, function(x) length(which(x >
        0))), Neg.interactions = apply(moose.bootstrap.spatial$direct_coef_means[1:42,
        2:43], 1, function(x) length(which(x <
        0))), row.names = NULL) %>% dplyr::mutate(OTU = gsub("X",
    "", OTU)) %>% left_join(taxonomy) %>%
    dplyr::arrange(Group)
knitr::kable(summary_table, caption = "Summary statistics of OTU co-occurrences")
```

Table 1: Summary statistics of OTU co-occurrences

| OTU | Ave.occurrence | Pos.interactions | Neg.interactions | Group |
|---|---|---|---|---|
| 338145 | 0.2043 | 6 | 4 | Actinobacteria:Coriobacteriia |
| NewReferenceOTU314 | 0.6637 | 2 | 1 | Actinobacteria:Coriobacteriia |
| NewCleanUpReferenceOTU69 | 0.6832 | 2 | 1 | Actinobacteria:Coriobacteriia |
| NewCleanUpReferenceOTU74 | 0.7562 | 3 | 2 | Actinobacteria:Coriobacteriia |
| 206930 | 0.4799 | 4 | 4 | Cyanobacteria:4C0d-2 |
| NewReferenceOTU123 | 0.5473 | 6 | 3 | Cyanobacteria:4C0d-2 |
| NewCleanUpReferenceOTU73 | 0.5461 | 0 | 2 | Cyanobacteria:Chloroplast |
| 322906 | 0.9437 | 3 | 4 | Firmicutes:Clostridia |
| 325706 | 0.6148 | 0 | 1 | Firmicutes:Clostridia |
| 4317006 | 0.0372 | 5 | 4 | Firmicutes:Clostridia |
| 4480841 | 0.5291 | 2 | 1 | Firmicutes:Clostridia |
| 4314603 | 0.4536 | 2 | 1 | Firmicutes:Clostridia |
| 4417708 | 0.0843 | 7 | 4 | Firmicutes:Clostridia |
| 296918 | 0.7692 | 1 | 4 | Firmicutes:Clostridia |
| 266952 | 0.6309 | 3 | 3 | Firmicutes:Clostridia |
| 294064 | 0.5084 | 5 | 3 | Firmicutes:Clostridia |
| 294262 | 0.2403 | 6 | 1 | Firmicutes:Clostridia |
| 579159 | 0.2068 | 3 | 2 | Firmicutes:Clostridia |
| 333577 | 0.3276 | 2 | 1 | Firmicutes:Clostridia |
| 1038874 | 0.3899 | 2 | 0 | Firmicutes:Clostridia |

| OTU | Ave.occurrence | Pos.interactions | Neg.interactions | Group |
|---|---|---|---|---|
| 129755 | 0.3879 | 3 | 0 | Firmicutes:Clostridia |
| 738351 | 0.6364 | 1 | 0 | Firmicutes:Clostridia |
| 4295783 | 0.3693 | 1 | 0 | Firmicutes:Clostridia |
| 574585 | 0.6326 | 1 | 5 | Firmicutes:Clostridia |
| NewReferenceOTU108 | 0.0809 | 8 | 3 | Firmicutes:Clostridia |
| NewReferenceOTU74 | 0.3172 | 1 | 0 | Firmicutes:Clostridia |
| NewReferenceOTU76 | 0.2891 | 5 | 6 | Firmicutes:Clostridia |
| NewCleanUpReferenceOTU76 | 0.0827 | 6 | 4 | Firmicutes:Clostridia |
| NewCleanUpReferenceOTU157 | 0.0899 | 3 | 0 | Firmicutes:Clostridia |
| NewCleanUpReferenceOTU289 | 0.1515 | 5 | 4 | Firmicutes:Clostridia |
| 4396877 | 0.0928 | 5 | 1 | Firmicutes:Erysipelotrichi |
| NewCleanUpReferenceOTU42 | 0.5286 | 3 | 4 | Firmicutes:Erysipelotrichi |
| 339838 | 0.0630 | 1 | 1 | Tenericutes:Mollicutes |
| 513605 | 0.0299 | 7 | 1 | Tenericutes:Mollicutes |
| 4446732 | 0.6627 | 4 | 2 | Tenericutes:Mollicutes |
| 1108356 | 0.5123 | 4 | 2 | Tenericutes:Mollicutes |
| 921020 | 0.1596 | 5 | 1 | Tenericutes:Mollicutes |
| NewReferenceOTU413 | 0.2271 | 5 | 2 | Tenericutes:Mollicutes |
| NewReferenceOTU493 | 0.2445 | 4 | 1 | Tenericutes:Mollicutes |
| NewReferenceOTU390 | 0.6923 | 0 | 0 | Tenericutes:Mollicutes |
| NewReferenceOTU404 | 0.1345 | 8 | 2 | Tenericutes:Mollicutes |
| NewReferenceOTU528 | 0.4842 | 4 | 4 | Tenericutes:RF3 |

For making plots, it is useful to extract an adjacency matrix of predicted interactions so that we can use `igraph` to construct informative networks

```
interactions <- moose.bootstrap.spatial$direct_coef_means[1:42,
    2:43]

# Change names of OTUs to match the
# summary table format
rownames(interactions) <- gsub("X", "", rownames(interactions))
colnames(interactions) <- gsub("X", "", colnames(interactions))

# Create an adjacency matrix
adj.mat <- igraph::graph.adjacency(interactions,
    weighted = T, mode = "undirected")

# Delete edges that represent weak
# interactions
adj.mat <- igraph::delete.edges(adj.mat,
    which(abs(igraph::E(adj.mat)$weight) <=
        0.1))
cols <- c(grDevices::adjustcolor("blue",
    alpha.f = 0.5), grDevices::adjustcolor("red4",
    alpha.f = 0.5))
igraph::E(adj.mat)$color <- ifelse(igraph::E(adj.mat)$weight <
    0, cols[1], cols[2])

# Add colours to show Groups
cols <- c("#000000", "#009E73", "#e79f00",
    "#9ad0f3", "#0072B2", "#D55E00", "#CC79A7")
```

```
vertex_groups <- summary_table$Group[which(summary_table$OTU %in%
    igraph::V(adj.mat)$name)]
igraph::V(adj.mat)$color <- cols[as.factor(vertex_groups)]
igraph::V(adj.mat)$label <- ""

# Adjust weights for easier visualisation
# of interaction strengths
igraph::E(adj.mat)$width <- abs(igraph::E(adj.mat)$weight) *
    1.2

# Remove vertices with no edges (no
# interactions)
adj.mat <- igraph::delete.vertices(adj.mat,
    igraph::degree(adj.mat) == 0)
par(mar = c(0, 0, 0, 10))
set.seed(25)
plot(adj.mat, edge.curved = 0.2, vertex.size = 6.6,
    vertex.frame.color = grDevices::adjustcolor("black",
        alpha.f = 0.8))
legend("bottomright", bty = "n", legend = levels(as.factor(vertex_groups)),
    col = cols, border = NA, pch = 16, cex = 0.9,
    inset = c(-0.45, 0))
```

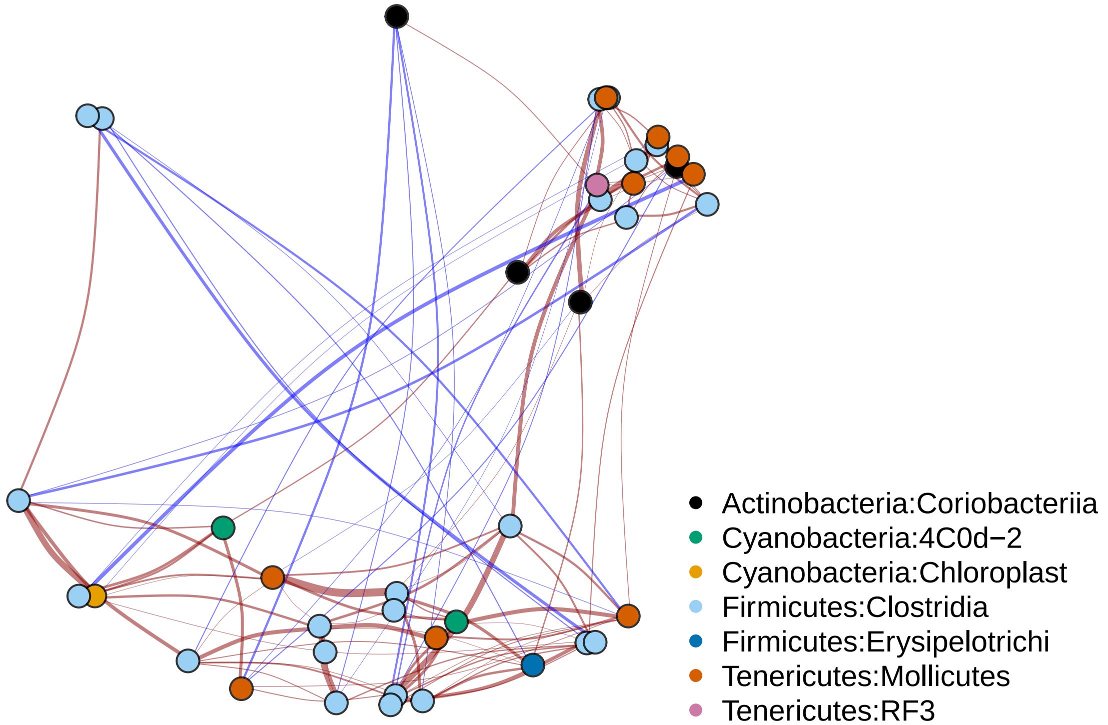
